# IL-17A Restrains Antiviral Immunity to Promote Chikungunya Virus Infection and Pathogenesis in the Heart

**DOI:** 10.64898/2026.08.07.743518

**Authors:** Shazeed-Ul Karim, Prince M.D. Denyoh, Sabin Shrestha, Ayokanmi Osobukola, Nathan S. Bai, Fengwei Bai

## Abstract

Chikungunya virus (CHIKV) infection is increasingly linked to cardiovascular complications, but the mechanisms underlying CHIKV-induced cardiovascular disease (CVD) remain unclear, and targeted therapies are lacking. Although elevated interleukin-17A (IL-17A) levels have been reported in CHIKV patients and associated with cardiovascular pathology, its role in CHIKV-induced cardiac disease is poorly defined. To address this question, we employed our newly developed heterozygous interferon α/β/γ receptor-deficient (*Ifnag*^+/−^) mice and primary human cardiac fibroblasts to investigate the contribution of IL-17A signaling to CHIKV-associated cardiac pathology. We found that CHIKV infection induced IL-17A production in the heart, and that mice deficient in *Il17a* (*Il17a^−/−^*) and in its receptor gene, *Il-17ra* (*Il17ra^−/−^*), exhibited marked resistance to CHIKV infection in both cardiac tissue and primary cardiac fibroblasts. Genetic deletion of IL-17A signaling significantly enhanced type I interferon responses and decreased viral burden in mouse hearts. Interestingly, blockade of IL-17RA with an FDA-approved monoclonal antibody for plaque psoriasis, Brodalumab, drastically increased type I interferon production and reduced viral replication in both human cardiac fibroblasts and human embryonic kidney 293 (HEK 293) cells. In addition, inhibition of IL-17A signaling suppressed the expression of pro-inflammatory mediators, including *Il-1β*, *Tnf-α*, and *Cxcl2*, reduced immune cell infiltration into cardiac tissue, and mitigated cardiac injury. Importantly, therapeutic blockade of IL-17A signaling after CHIKV infection reduced viral replication in both the heart and circulation. Collectively, these findings identify IL-17A signaling as a critical regulator of CHIKV replication and cardiac inflammation and highlight the IL-17A/IL-17RA axis as a promising therapeutic target for CHIKV-associated cardiovascular disease.

**Importance:** Chikungunya virus (CHIKV) infection has been frequently associated with cardiovascular complications, yet the host pathways that promote viral infection and cardiac injury remain poorly understood. Here, we identify IL-17A signaling as a previously unrecognized regulator of CHIKV pathogenesis in the heart. Using a novel heterozygous interferon receptor-deficient mouse model and primary human cardiac fibroblasts, we demonstrate that IL-17A signaling facilitates CHIKV replication via suppressing antiviral type I interferon responses. Genetic deletion or pharmacological blockade with an FDA-approved monoclonal antibody of IL-17A signaling reduced viral burden, attenuated inflammatory cytokine production, limited immune cell infiltration, and protected against cardiac injury. Importantly, therapeutic inhibition of IL-17A signaling after infection remained effective in reducing viral replication in both cardiac tissue and circulation and mitigating cardiac damage. These findings reveal a critical role for the IL-17A/IL-17RA axis in linking antiviral immunity to CHIKV-induced cardiovascular disease and identify a potential translatable therapeutic target for CHIKV-caused cardiac complications.

## Introduction

Chikungunya virus (CHIKV) is a single-stranded, positive-sense RNA virus belonging to the genus *Alphavirus* in the family *Togaviridae*. CHIKV is transmitted by the bite of an infected mosquito (*Aedes aegypti* and *A. albopictus*). Historically restricted to tropical regions, CHIKV now poses an increasing threat due to climate-driven northward expansion of the mosquito vectors, with documented local transmission in multiple U.S. states(1–3). Although the typical symptoms of chikungunya disease are acute fever and chronic arthritis, there are many clinical reports that detail a wide range of cardiac symptoms manifested during CHIKV infection, including arrhythmias, palpitations, abnormal electrocardiograms and echocardiograms, myocardial infarctions, heart failure, cardiac arrest, cardiomegaly, and myocarditis(4, 5). While some of these manifestations may be due to immune responses to the infection, including the effects of cytokines and inflammation, other symptoms result from direct viral invasion of cardiac tissue(5–8). The most prevalent cardiovascular disease (CVD) complication of CHIKV infection, which is found across nearly all age groups, is myocarditis, which results when the virus directly infects the cardiac tissues and causes damage(5, 7, 9). Supporting this notion, all currently identified CHIKV receptors, including prohibitin (PHB), matrix remodeling-associated protein 8 (MXRA8), and CD147, are expressed on the surface of human cardiomyocytes(8). Furthermore, CHIKV antigens have been detected in cardiac tissues, and elevated viral titers are associated with more severe cardiac injury(10–13).

Interleukin-17A (IL-17A) mediates a wide array of immunological processes, ranging from host defense and tissue repair to the progression of cancer and autoimmune disorders(14). Clinical studies have frequently suggested a link between elevated IL-17A levels in patients’ serum and the pathogenesis of CHIKV(15, 16). Elevated levels of IL-17A are crucial for the development of inflammatory arthritis and various allergic conditions (17, 18). Clinical studies have also reported elevated IL-17A levels in patients post-CHIKV infection, and growing evidence links IL-17A to cardiovascular pathologies, including heart failure(19). Our previous investigations found that IL-17A-deficient (*Il17a^−/−^)* and IL-17A receptor-deficient (*Il17ra^−/−^)* mice exhibit reduced viremia and CHIKV in the footpad following CHIKV infection(20). In addition, our study suggests that *Il-17a* is highly expressed in the hearts of CHIKV-infected mice compared with uninfected control mice, and that CHIKV RNA is almost undetectable in the hearts of *Il17a^−/−^* mice inoculated with CHIKV. These findings support the hypothesis that IL-17A signaling plays a crucial role in CHIKV pathogenesis; however, the mechanism by which IL-17A mediates CHIKV-induced CVD remains to be characterized.

CHIKV infection induces the production of proinflammatory cytokines, chemokines, and type I interferons (IFN-α/β)(21). IFNα/β play a key role in limiting CHIKV replication in both human and mouse models(22). We recently developed and validated heterozygous interferon receptor-deficient (*Ifnar1^+/−^* and *Ifnag^+/−^*) mouse models of CHIKV infection in the heart, demonstrating viral cardiac tropism and immune-mediated myocardial injury(23). Importantly, our mouse models permit viral replication in cardiac tissues while preserving sufficient immune competence to interrogate immunoregulatory mechanisms(23). Therefore, in this study, we used *Ifnag^+/−^* mice to investigate the role of IL-17A signaling in the pathogenesis of CHIKV-induced cardiac diseases, and our results suggest that targeting the IL-17A/IL-17RA axis may represent a promising therapeutic strategy for CHIKV-induced CVDs.

## Results

### CHIKV induces IL-17A production, which facilitates CHIKV infection in the heart

To determine whether IL-17A is induced in the hearts of mice, we compared *Il17a* transcript levels in *Ifnag^+/−^* mice after CHIKV infection via footpad, and observed upregulation in infected hearts relative to uninfected controls (Fig. 1A), consistent with our previous findings in footpad tissues (20). To determine whether IL-17A signaling similarly influences viral dissemination to the heart, we infected WT, *Il17a^−/−^*, *Il17ra^−/−^*, and *Ifnag^+/−^* mice via the footpad. On day (D) 1 post-infection (p.i.), cardiac tissues were collected after perfusion, and CHIKV *E1* RNA was quantified by RT-qPCR targeting the CHIKV *E1* gene, with normalization to cellular *β-actin*. We found that both *Il17a^−/−^*and *Il17ra^−/−^* mice exhibited a significantly reduced CHIKV burden compared to WT and *Ifnag^+/−^*mice (Fig. 1B). As expected, *Ifnag^+/−^* mice displayed higher viral loads than WT mice, *Il17a^−/−^*, and *Il17ra^−/−^* mice, suggesting that IL-17A signaling facilitates CHIKV infection in the heart (Fig. 1B). We collected cardiac tissues on D1 p.i. because our recent study showed that CHIKV burden peaks in the heart of heterozygous *Ifnar1^+/−^* and *Ifnag^+/−^* mice on D1, followed by a rapid decline from D2 to D4, and then became very low, even undetectable, by D5 p.i.(23).

**Figure 1.**
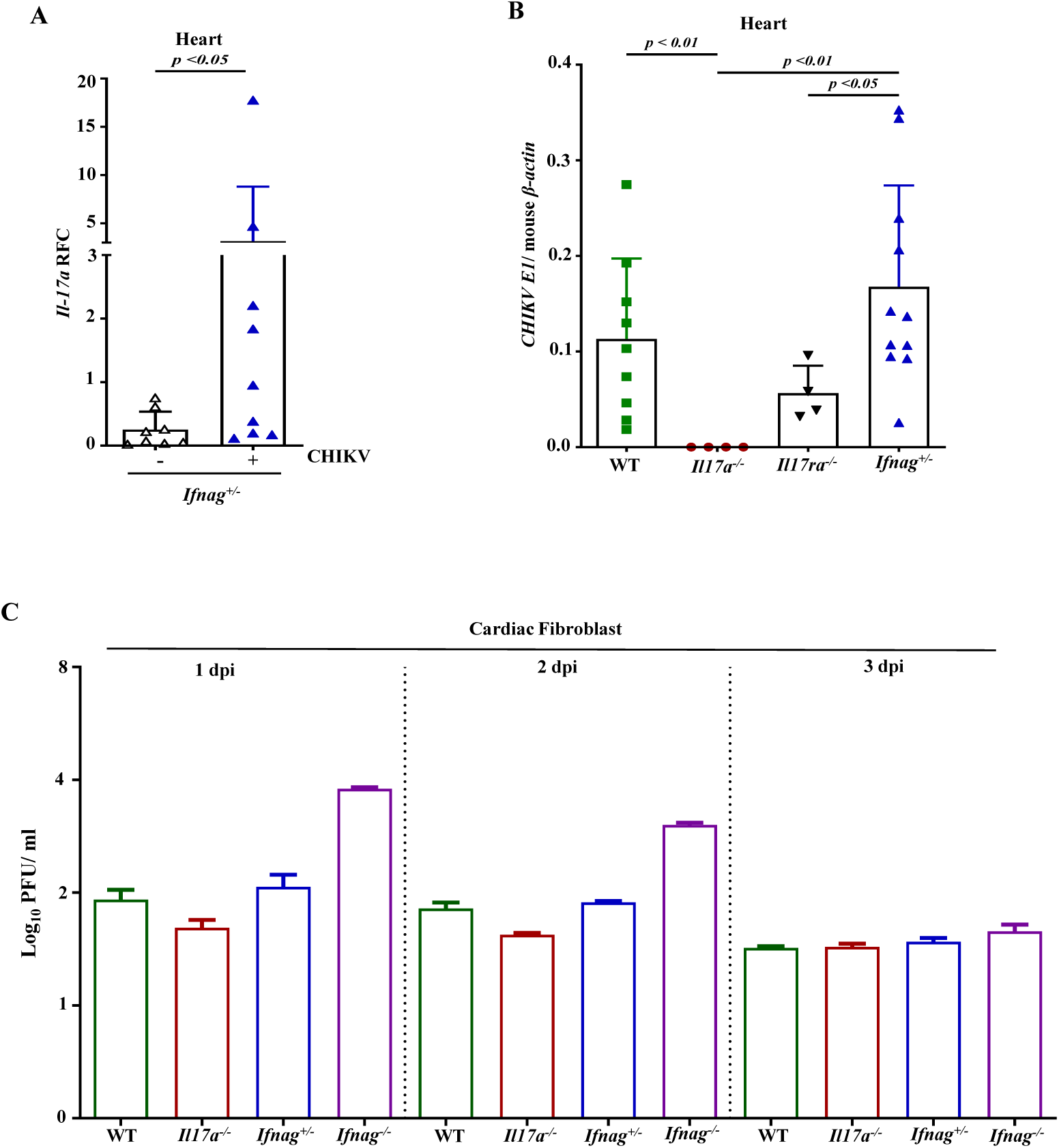
IL-17A signaling facilitates CHIKV infection in the heart. (A) *Ifnag^+/−^*mice were infected with CHIKV, and *Il17a* expression in the heart on D1 post-infection (p.i) was quantified by RT-qPCR and compared with uninfected controls. (B) *CHIKV E1* RNA levels in the heart of WT, *Il17a^−/−^*, *Il17ra^−/−^*, and *Ifnag^+/−^* mice *on* D1 p.i.. (C) Primary cardiac fibroblasts of WT, *Il17a^−/−^*, *Ifnag^+/−^*, and *Ifnag^−/−^* mice were infected with CHIKV (*n =*2). Viral titers in the media were quantified by plaque-forming assay at 1, 2, and 3 days post-infection. The statistical significance in (A) and (B) was determined using the Mann-Whitney U test.

Recent studies have suggested that cardiac fibroblasts are the primary cellular targets for CHIKV among various cardiac populations, including cardiac endothelial cells, cardiac muscle cells, and leukocytes(24, 25). Although representing a smaller proportion of cardiac cells than cardiomyocytes, cardiac fibroblasts and endothelial cells are more susceptible to alphavirus infection because they express high levels of MXRA8, the major entry receptor for alphaviruses, including CHIKV (26–28). This led us to evaluate whether CHIKV infection in cardiac fibroblasts is regulated by IL-17A signaling. Primary cardiac fibroblasts were isolated from WT, *Il17a^−/−^*, *Ifnag^+/−^* and *Ifnag^−/−^*mice, cultured at the same concentration, and infected with CHIKV at an MOI of 1.0 *in vitro*. Culture supernatants were collected at D1 p.i., and viral titers were determined by plaque-forming assay. Cardiac fibroblasts isolated from *Il17a^−/−^* mice exhibited relatively reduced susceptibility to CHIKV infection compared to those cells from WT, *Ifnag^+/−^*, and *Ifnag^−/−^* mice, with the cardiac fibroblasts from *Ifnag^−/−^* mice exhibiting the highest viral burden (Fig. 1C). These results suggest that IL-17A signaling promotes CHIKV infection in cardiac fibroblasts. In summary, our *in vivo* and *in vitro* results demonstrate that IL-17A signaling promotes CHIKV replication in cardiac tissues.

### IL-17A signaling blockage inhibits CHIKV infection in the cardiac tissues

To determine if blocking IL-17A signaling reduces CHIKV infection in the heart, three-week-old *ifnag^+/−^* mice were administered either an anti-IL-17A monoclonal antibody (α-IL17A, 100 μg/mouse) or an isotype control antibody (IgG, 100 μg/mouse) via intraperitoneal (i.p.) injection. In the pre-treatment experiments, antibodies were administered on D-1 and D0, and CHIKV was injected via footpad inoculation on D0. On D1 p.i., blood samples were collected, and the mice were anesthetized and perfused to collect the heart for viral quantification (Fig. 2A). Our results suggest that α-IL-17A treatment significantly reduced CHIKV infection in the heart as measured by RT-qPCR (Fig. 2B) and plaque-forming assay (Fig. 2C), compared to isotype control mice. Consistently, α-IL17A -treated mice also exhibited lower CHIKV RNA in the blood (Fig. 2D). To further characterize the spatial distribution of CHIKV and confirm the protective effects of IL-17A neutralization, we performed immunohistochemistry (IHC) on cardiac sections. Our previous results suggested that the left atrium and left ventricle exhibited higher viral loads than the right atrium and right ventricle in hearts with CHIKV infection(23). Therefore, we evaluated the presence of the CHIKV envelope protein 2 (E2) in longitudinal heart sections from *ifnag^+/−^* mice that received either α-IL-17A or an isotype control Ab and were then infected with CHIKV. The IHC results showed that α-IL-17A -treated mice exhibited a reduced number of CHIKV E2-positive cells compared to the isotype Ab-treated control group in the left atrium (Fig. 2E) and the left ventricle (Fig. 2G) on D1 p.i. These results further confirm that blocking IL-17A signaling effectively limits CHIKV replication in cardiac tissues. Quantification of CHIKV E2-positive cells, expressed as the percentage of CHIKV E2-positive cells relative to the total number of DAPI-positive nuclei, further demonstrated a significant reduction in CHIKV E2-positive cells in the α-IL-17A treated group in the left atrium (Fig. 2F) and left ventricle (Fig. 2H). Collectively, these findings provide additional evidence that IL-17A signaling blockage limits CHIKV burden in cardiac tissues.

**Figure 2.**
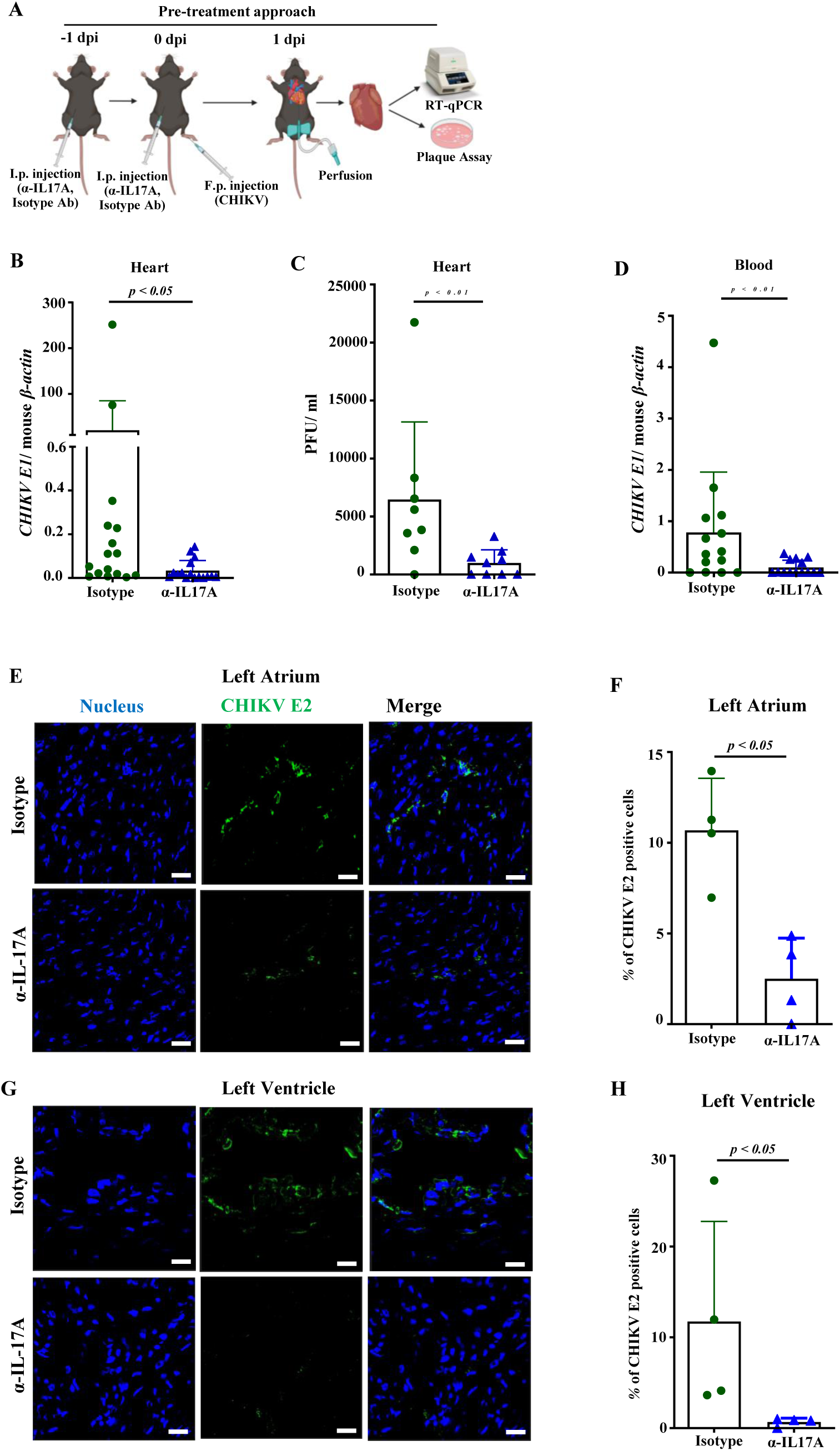
Blockage of IL-17A signaling inhibits CHIKV infection in cardiac tissues. (A) Experimental schematic showing *Ifnag^+/−^* mice treated with α-IL-17A antibody or isotype control antibody on D-1 (pre-treatment) and D0 via intraperitoneal (i.p.), followed by CHIKV infection on D0 via footpad (FT). Hearts were collected at D1 p.i. after perfusion for viral quantification. (B) *CHIKV E1* RNA levels in the heart at D 1 were measured by RT-qPCR (normalized to mouse *β-actin*). (C) Infectious CHIKV titers in the heart were measured by plaque-forming assay. (D) *CHIKV E1* RNA in the blood on D 1 p.i. was quantified by RT-qPCR. (E, G) Representative immunohistochemistry (IHC) images showing CHIKV envelope protein 2 (E2) presence in the left atrium (E) and left ventricle (G) of longitudinal heart sections collected at D1 p.i.. (F, H) Quantification of CHIKV E2-positive cells, expressed as the percentage of CHIKV E2-positive cells relative to the total number of DAPI-positive nuclei in the left atrium (F) and left ventricle (H). The statistical significance was determined using Mann-Whitney U tests. Images were acquired using a Stellaris STED confocal microscope (Leica) at 63× magnification. The scale bars represent 20µm for all images. Fig. 2A was created with BioRender.

### Blockage of IL-17A signaling enhances interferon response and suppresses pro-inflammatory cytokine expression in the heart

Interferons (IFNs) are pivotal components of innate immunity and are essential for host defense against viral pathogens. Our previous findings demonstrated that IL-17A facilitates CHIKV infection by inhibiting IFN-α2 expression through modulation of Interferon Regulatory Factor-5 (IRF-5), IRF-7, IFN-stimulated gene 49, and Mx1 (20). Therefore, we investigated whether neutralization of IL-17A could restore the IFN response in cardiac tissues. Similarly, *Ifnag^+/−^* mice were treated with either an α-IL-17A or an isotype control on D-1 and D0, then followed by CHIKV infection. The hearts were isolated after perfusion on D1 for RT-qPCR analysis, and the results indicate a trend of increasing expression of type I and type II interferons, including *Ifn-α*, *Ifn-β*, and *Ifn-γ*, in the α-IL-17A -treated mice compared to the isotype control Ab-treated mice, with *Ifn-α* showing a statistically significant increase (Fig. 3 A-C). These data suggest that blocking IL-17A signaling leads to a more robust type I interferon response, thereby facilitating inhibition of CHIKV infection in the heart.

**Figure 3.**
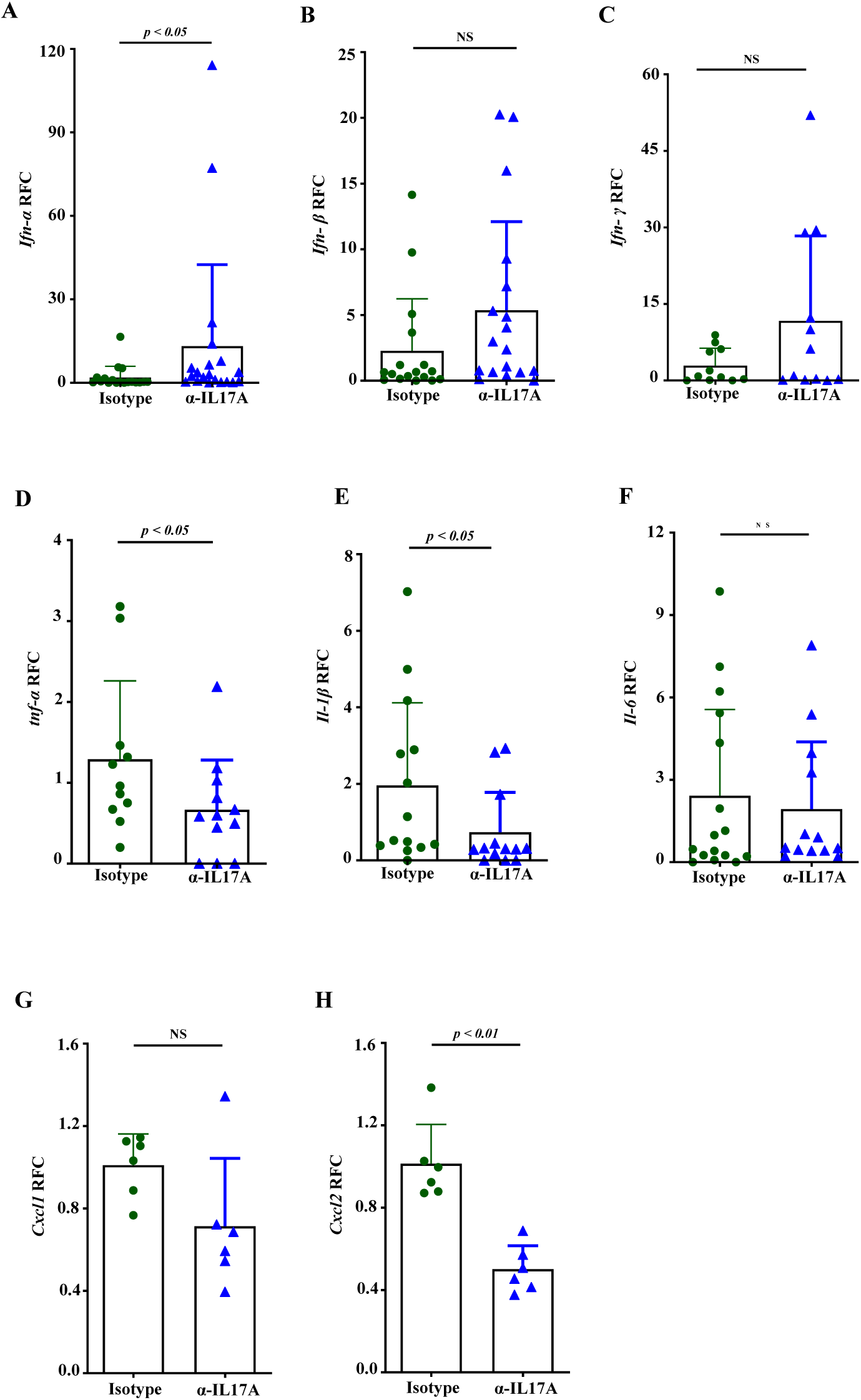
Blockage of IL-17A signaling enhances interferon responses and suppresses pro-inflammatory cytokine and chemokine expression in the heart. *Ifnag^+/−^* mice were treated with either α-IL-17A antibody or isotype control on D-1 (pre-treatment) and D0, followed by CHIKV infection, and hearts were collected on D1 p.i. after perfusion. RT-qPCR analysis was performed to quantify the expression of interferons (A-C), including *Ifn-α*, *Ifn-β*, and *Ifn-γ*; pro-inflammatory cytokines (D-F), including *Il-6*, *Il-1β*, and *Tnf-α*; and chemokines (G and H), including *Cxcl1* and *Cxcl2*. The statistical significance was determined using Mann-Whitney U tests.

Viral recognition by host pattern recognition receptors (PRRs) typically triggers the expression of pro-inflammatory cytokines and chemokines, thereby recruiting immune cells to the site of infection(29–31). However, excessive cytokine levels can lead to collateral damage in cardiac tissue, including apoptosis, reduced contractility, and cardiomyocyte hypertrophy(32, 33). We assessed the expression levels of key pro-inflammatory markers, including interleukin-6 (*Il-6*), interleukin-1β (*Il-1β*), tumor necrosis factor (*Tnf-α*), and the chemokines, including C-X-C motif chemokine ligand (*Cxcl*)1 and *Cxcl2*, in the hearts of CHIKV-infected *Ifnag^+/−^* mice (Fig. 3D-H). The isotype control group exhibited significantly elevated cytokines and chemokines, including *Tnf-α* (Fig. 3D), *Il-1β* (Fig. 3E), and *Cxcl2* (Fig. 3H). These elevated levels of cytokines and chemokines in the isotype Ab-treated group are consistent with the disease severity associated with CHIKV pathogenesis reported in previous studies (34, 35). In contrast, mice treated with the α-IL-17A showed significantly reduced expression of these pro-inflammatory mediators (Fig 3D-H). These results indicate that neutralizing IL-17A signaling augments antiviral IFN defenses, thus inhibiting CHIKV replication and indirectly limiting the potentially pathological cytokines and chemokines in the heart.

### IL-17A facilitates immune cell infiltration and cardiac tissue pathology

After cardiac infection, infiltrating innate immune cells, such as macrophages and neutrophils, release pro-inflammatory cytokines, reactive oxygen species, and proteolytic enzymes that injure cardiomyocytes and amplify local inflammation(36, 37). In addition, activated T lymphocytes play important roles in the development of myocarditis and other myocardial diseases by recognizing cardiac antigens or cross-reactive microbial antigens, directly killing cardiomyocytes and sustaining myocardial inflammation even after the initial pathogen burden declines(36, 38). Persistent immune cell infiltration promotes fibroblast activation and extracellular matrix remodeling, leading to fibrosis, impaired cardiac function, and progression to inflammatory cardiomyopathy or heart failure(39). To quantify cardiac immune cell infiltration, we performed flow cytometry on single-cell suspensions from the hearts of CHIKV-infected *Ifnag^+/−^*mice treated with α-IL-17A or isotype control on D1 p.i.. The results showed that IL-17A blockage significantly reduced leukocyte recruitment to the heart, decreasing total CD45⁺ cells from 2.03% in isotype controls to 0.84% in α-IL-17A-treated mice, as shown in the representative flow cytometry dot plots (Fig. 4A). Consistently, both representative histograms and bar graph confirmed a significant reduction in overall leukocyte infiltration following IL-17A blockage (Fig. 4A). Of the total leukocytes, neutrophil infiltration was markedly reduced by α-IL-17A treatment, declining from 22.6% in isotype controls to 7.83% (Fig. 4B). Likewise, T-cell recruitment was also significantly decreased, with T cells comprising 17.4% in isotype controls versus 9.10% in the heart of α-IL-17A -treated mice (Fig. 4C). Together, these findings identify IL-17A as a key driver of early inflammatory cell infiltration into the heart during CHIKV infection.

**Figure 4.**
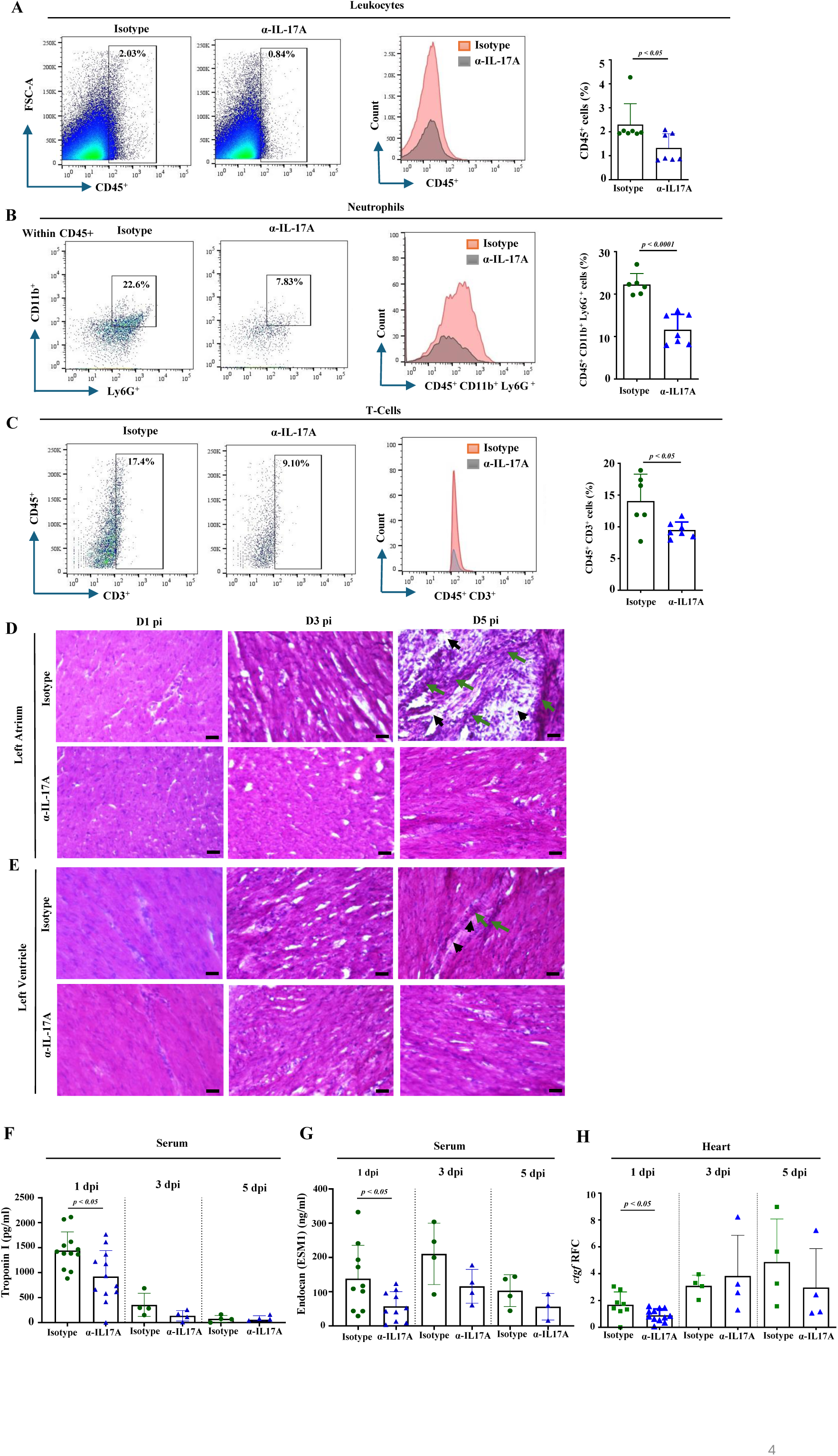
IL-17A signaling blockage reduces immune cell infiltration and tissue injury in the heart. *Ifnag^+/−^* mice were treated with either α-IL-17A antibody or isotype control at D −1 and D0 p.i., followed by CHIKV infection. Heart tissues and blood samples were collected at D1, D3, and D5 p.i., Blood samples were allowed to clot and subsequently centrifuged to obtain serum. (A-C) Representative flow cytometry dot plots showing the percentages of immune cell populations and histograms showing immune cell counts in the heart of isotype Ab-treated and α-IL-17A-treated mice. Corresponding bar graphs summarize the percentages of each immune cell population. Populations are defined as follows: (A) leukocytes as CD45⁺; (B) neutrophils as CD45⁺ CD11b⁺ Ly6G⁺; and (C) T cells as CD45⁺ CD3⁺. Sample sizes range from n = 6 to 7 per group. (D and E) Histopathological assessment of cardiac tissue at D1, D3, and D5 p.i. using Hematoxylin and Eosin (H&E) staining. Representative images of the left atrium (D) and left ventricle (E) illustrate immune cell infiltration (green arrows) and tissue damage (black arrows) in isotype versus α-IL-17A-treated hearts. The images were captured by the Nikon Eclipse 80i microscope (Nikon, Japan) at 40× magnification. The scale bars represent 200 µm for all images. Serum levels of the cardiac biomarkers Troponin I (F) and Endocan (G) were quantified by ELISA. (H) The expression of cardiac connective tissue growth factor (*Ctgf*) was analyzed by RT-qPCR. The statistical significance (A-C and F-H) was determined using an unpaired two-tailed Student’s *t*-test.

To assess the pathologic impact of the inflammatory responses, we performed histological analysis using Hematoxylin and Eosin (H&E) staining. In the isotype Ab-treated *Ifnag^+/−^* mice, the myocardium, particularly within the left atrium, was characterized by dense immune cell infiltration (green arrows), and apparent structural damage (black arrows) at D5 p.i., indicating the development of myocarditis (Fig. 4D). While the pathology was most pronounced in the left atrium, the left ventricle of isotype Ab-treated mice also exhibited evidence of immune cell infiltration and myocardial damage, albeit to a lesser extent than the atrial tissues (Fig. 4E). These observations further suggest that IL-17A signaling is a critical mediator of CHIKV-induced cardiac pathology.

Recent clinical studies have shown that cardiac injury during severe CHIKV infection is associated with elevated cardiovascular biomarkers. Fatal cases exhibit markedly increased levels of Endocan-1 (endothelial cell-specific molecule-1, ESM-1) and higher Troponin I concentrations, indicating that endothelial dysfunction and direct myocardial injury contribute to disease severity(40). In addition, connective tissue growth factor (CTGF), a sensitive marker of cardiac fibrosis, provides a more accurate measure of early fibrotic remodeling at the transcript level than conventional histological assessment(41). Elevated levels of these biomarkers are therefore indicative of severe cardiac damage. To further confirm that IL-17A signaling contributes to cardiac injury, serum and heart tissues were collected from CHIKV-infected *Ifnag^+/−^* mice with either α-IL-17A or an isotype control Ab and analyzed at D1, D3, and D5 p.i.. ELISA results revealed significantly lower serum levels of Troponin I and Endocan in α-IL-17A -treated mice on D1 p.i., with this reduction remaining evident at subsequent time points (Fig. 4F and G). Consistently, RT-qPCR analysis showed reduced *Ctgf* expression in α-IL-17A-treated mice compared with the control mice on D1 p.i. (Fig. 4H). Collectively, these findings indicate that IL-17A signaling promotes immune cell infiltration and myocardial injury, as well as potential fibrotic responses during CHIKV infection, whereas its blockade mitigates cardiac pathology.

### Therapeutic IL-17A neutralization restricts CHIKV burden in the heart and blood

To further evaluate clinical relevance and therapeutic potency, we infected *Ifnag^+/−^* mice with CHIKV and then treated them with α-IL17A or isotype control Ab at 6 hours p.i. (Fig. 5A). At D1 p.i., blood samples and hearts were collected after perfusion. Similar to the pre-treatment results, the therapeutic-treatment results showed a significant reduction of *CHIKV E1* RNA levels (Fig. 5B) and infectious titers (Fig. 5C) in the heart, as well as reduced viral RNA in the blood (Fig. 5D) in α-IL17A -treated mice compared to control mice.

**Figure 5.**
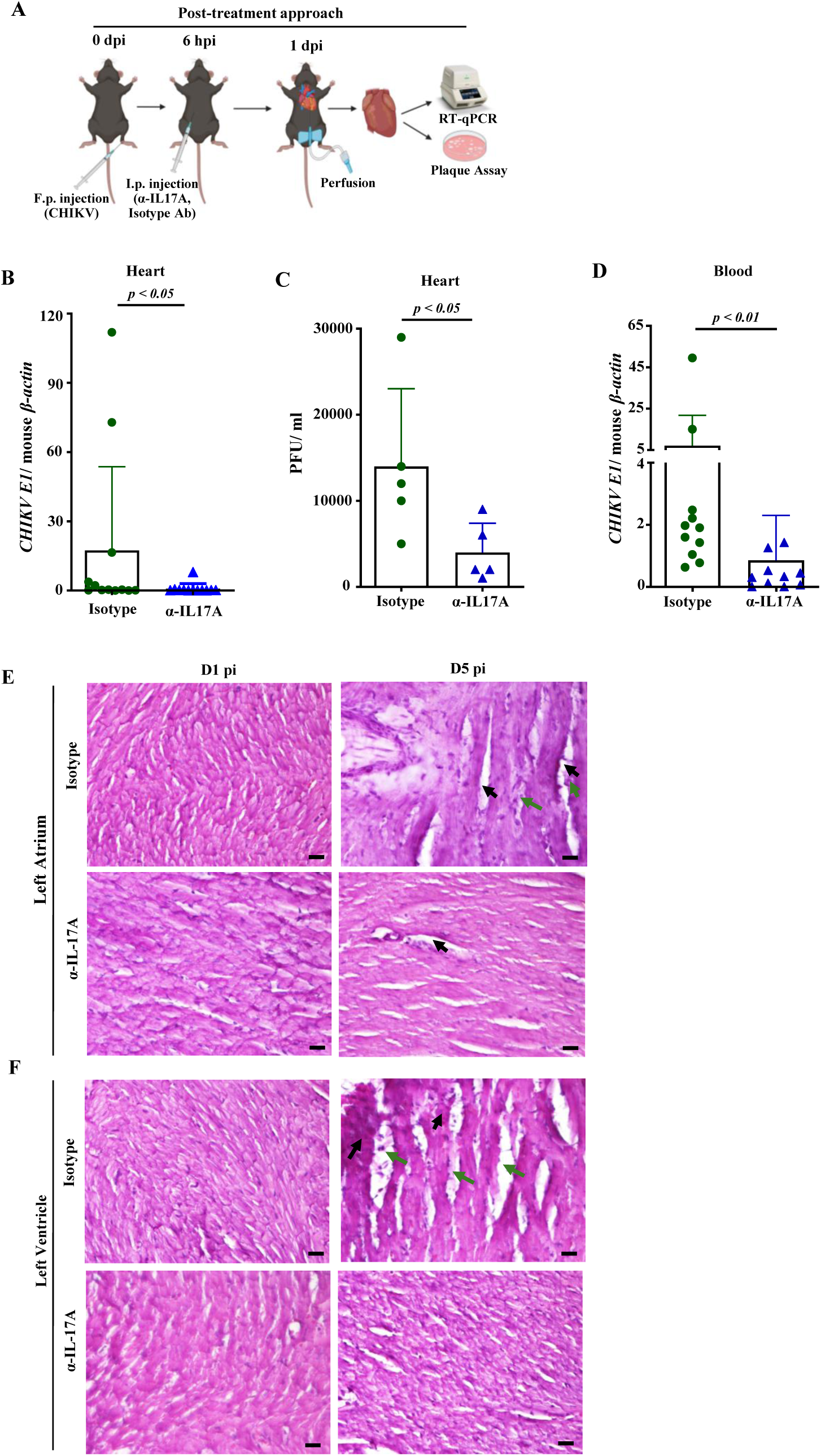
Therapeutic neutralization of IL-17A signaling restricts cardiac and systemic CHIKV burden. (A) Experimental schematic showing CHIKV-infected *Ifnag^+/−^* mice treated with α-IL-17A or isotype control Ab at 6 hours post-infection p.i. (Created with BioRender). Hearts were collected at D1 p.i. after perfusion for viral burden quantification. (A) *CHIKV E1* RNA levels were measured by RT-qPCR (normalized to mouse *β-actin*). (C) Infectious CHIKV titers were measured by plaque-forming assay. (D) *CHIKV E1* RNA in the blood on D1 p.i. was quantified by RT-qPCR. (E and F) Histopathological assessment of cardiac tissue at D1 and D5 p.i. using H&E staining. Representative images of the left atrium (E) and left ventricle (F) illustrate cell infiltration (green arrows) and tissue damage (black arrows) in isotype versus α-IL-17A-treated hearts. The images were captured by the Nikon Eclipse 80i microscope (Nikon, Japan) at 40× magnification. The scale bars represent 200 µm for all images. The statistical significance (B - D) was determined using the Mann-Whitney U test.

We further evaluated cardiac histopathology by H&E staining. At D1 p.i., neither α-IL17A-nor isotype-Ab-treated mice exhibited visible tissue damage in the left atrium or left ventricle (Fig. 6E and F). However, at D5 p.i., isotype-Ab-treated mice exhibited apparent cell infiltration (green arrows) and structural damage (black arrow) in both the left atrium and left ventricle compared to the α-IL17A-treated mouse group (Fig. 5E and F). These findings suggest that neutralization of IL-17A signaling, even post-CHIKV exposure, effectively restricts viral infection in the heart and circulation and reduces CHIKV-associated cardiac tissue damage, indicating the therapeutic potential of blocking IL-17A signaling to treat CHIKV-induced heart diseases.

**Figure 6.**
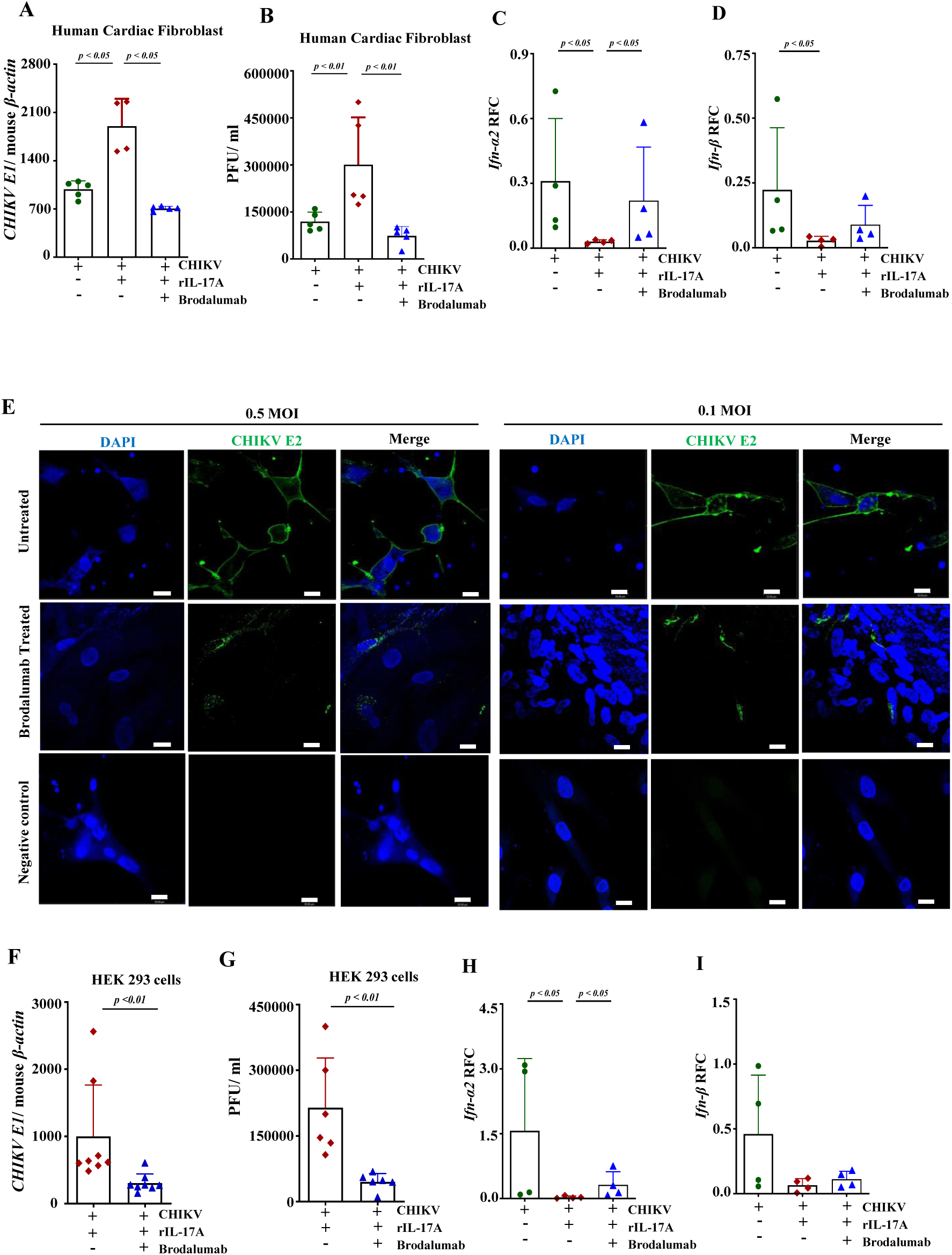
Blockage of IL-17A signaling reduces CHIKV replication in human primary cardiac fibroblasts and HEK 293 cells. Primary human cardiac fibroblasts (HCFs) were treated with Brodalumab (IL-17RA antagonist) for 3 hours prior to CHIKV infection and human recombinant IL-17A (rIL-17A) treatment. CHIKV infection was quantified by (A) RT-qPCR for *CHIKV E1* RNA and (B) plaque-forming assay for infectious titers in the medium. (C, D) The expression of human *Ifn-α2* and *Ifn-β* and were measured by RT-qPCR, (E) Representative immunofluorescence images of HCFs infected with CHIKV at MOI of 0.1 and 0.5. Cells were stained for CHIKV E2 protein (green) and DAPI for nuclei (blue). Images were acquired using a Stellaris STED confocal microscope (Leica) at 63× magnification. The scale bars represent 20µm. (F and G) HEK 293 cells were subjected to Brodalumab treatment prior to CHIKV infection and rIL-17A stimulation. Viral replication was determined by (F) RT-qPCR for *CHIKV E1* RNA and (G) plaque-forming assay. (H and I) The expression of human *Ifn-α2* and *Ifn-β* was measured by RT-qPCR. The statistical significance was determined using the Mann-Whitney U test.

### IL-17A signaling promotes CHIKV replication in human primary cardiac fibroblasts and HEK 293 cells

To extend our murine findings to human cells, we examined the role of IL-17A signaling in CHIKV replication using primary human cardiac fibroblasts (HCFs) and human embryonic kidney (HEK) 293 cells. Since these cells are not major producers of IL-17A, and to mimic the *in vivo* environment with IL-17A presence, we supplied recombinant human IL-17A (rIL-17A) to the cell culture. IL-17A signaling was antagonized by Brodalumab, an FDA-approved human monoclonal antibody targeting IL-17 receptor A (IL-17RA) for treating adult plaque psoriasis(42). HCFs were subjected to three conditions: CHIKV infection alone, CHIKV infection with rIL-17A, or Brodalumab pre-treatment followed by CHIKV infection and rIL-17A supplementation. Consistent with the results obtained in mouse cardiac fibroblasts (Fig. 1C) and cardiac tissues (Fig. 1B), IL-17A signaling enhanced CHIKV replication in primary human cardiac fibroblasts, whereas IL-17RA blockade with Brodalumab neutralized this effect (Fig. 6A and B). These results were further corroborated by immunocytochemical staining for CHIKV E2 protein. At MOIs of 0.1 and 0.5, Brodalumab treatment markedly reduced the number of CHIKV-positive cells compared with untreated controls, further confirming that IL-17A signaling augments CHIKV replication in both murine and human cardiac cells (Fig. 5E). In addition, consistent with murine results, human rIL-17A also inhibits type I interferon (α/β) expression, i.e., IFN-α2 and *Ifn-β*, whose effects can be reversed with Brodalumab (Fig. 6 C and D).

To determine whether these findings extend beyond cardiac cells, we infected human embryonic kidney (HEK) 293 cells with CHIKV in the presence or absence of Brodalumab. Consistent with results observed in primary human cardiac fibroblasts, Brodalumab pre-treatment significantly suppressed CHIKV replication, as evidenced by reduced viral RNA levels and plaque-forming units (PFUs) (Fig. 6F and G). As expected, rIL-17A treatment also reduced the expression of *Ifn-α2* and *Ifn-β* expression in CHIKV-infected HEK293 cells, whereas this effect was partially reversed following treatment with Brodalumab (Fig. 6H and I). Together, these findings demonstrate that the IL-17A/IL-17RA signaling axis promotes CHIKV replication in both primary human cardiac fibroblasts and HEK 293 cells, whereas pharmacological blockade of IL-17A signaling effectively inhibits viral infection, suggesting that targeting IL-17A signaling may be a promising therapeutic strategy for CHIKV-caused CVDs in humans.

## Discussion

IL-17A is the founding member of the IL-17 cytokine family, which also includes five other members, i.e., IL-17B to IL-17F (43, 44). IL-17A signaling is initiated when the cytokine binds to its receptor complex, primarily a heterodimer of IL-17RA and IL-17RC (45). The binding of IL-17A to its receptors recruits Act1, a multifunctional adapter protein that specifically binds to the receptor’s intracellular SEFIR domain(46). Act1 subsequently recruits various tumor necrosis factor receptor-associated factors (TRAFs), such as TRAF6, which activates the NF-*κ*B pathway and three mitogen-activated protein kinase (MAPK) pathways, i.e., JNK, p38, and ERK (47). These pathways collectively drive the transcription of inflammatory target genes. Additionally, IL-17A signaling regulates gene expression at the post-transcriptional level by controlling the stability of target mRNA through the formation of ribonucleoprotein (RNP) complexes that can either stabilize mRNA or accelerate its decay (47).

IL-17A is a pleiotropic cytokine that regulates a broad range of immune functions (48, 49). It is well recognized as a key mediator of inflammation (50) and plays important roles in the pathogenesis of allergic and autoimmune diseases, including rheumatoid arthritis (51–53), psoriasis (53, 54), asthma (55), and cancer(56). In contrast, the role of IL-17A during viral infections remains incompletely understood and appears to be highly context-dependent. Studies from our laboratory demonstrated that IL-17A promotes clearance of West Nile virus (WNV) from the brain by enhancing CD8^+^ T-cell cytotoxicity through activation of the PI3K-mTOR signaling pathway(57). Similarly, IL-17A has been shown to suppress Herpes Simplex Virus (HSV) infection by strengthening T-cell and Th1-mediated immune responses(58). However, IL-17A can also contribute to viral pathogenesis. For example, IL-17A exacerbates Coxsackievirus B3 infection by suppressing CD8^+^ T-cell activity and reducing IFN-γ production(59). Likewise, IL-17A promotes Theiler’s murine encephalomyelitis virus infection by inhibiting apoptosis in infected cells, thereby preventing their elimination(60).

Emerging evidence suggests that IL-17A also plays a pathogenic role during CHIKV infection. Clinical studies have reported elevated serum IL-17A levels in CHIKV-infected patients, which strongly correlate with arthritis severity and joint swelling, implicating IL-17A in the development of chronic inflammatory manifestations(34). Consistent with these observations, one of our previous studies identified a novel role for IL-17A in promoting CHIKV infection in both murine models and human cell lines(20). Mechanistically, IL-17A enhanced viral replication by suppressing IFN-α2 expression, a major subtype of IFN-α, and modulating the activity of key antiviral factors, including IRF-5, IRF-7, and ISG49(20). Consistent with our findings, a recent study with hepatitis B virus (HBV) infection found that IL-17A pretreatment attenuates the anti-HBV efficacy of interferon-α by reducing activation of the interferon-stimulated gene factor 3 (ISGF3) complex (phosphorylated (p)-signal transducer and activator of transcription (STAT1)/p-STAT2/IRF9) and antiviral-related ISGs (ISG15, ISG20, and Mx1) in hepatitis B virus-expressing HepG2 cells(61). In addition to its role in viral infections, accumulating evidence has linked IL-17A signaling to the pathogenesis of several CVDs, including atherosclerosis (62–64), hypertension(65), stroke(66), acute myocardial infarction(67), myocarditis (68–70), and heart failure(71). Despite these associations, the mechanisms by which IL-17A contributes to CHIKV-induced CVDs remain largely unknown.

Our recent study suggests that C57BL/6J wild-type (WT) mice are resistant to CHIKV infection and without causing significant damage in the heart, while homozygous *Ifnar1^−/−^* and *Ifnag^−/−^* mice (both in the C57BL/6J background) die of CHIKV infection within 3 days, limiting the use of these animals to study the disease mechanisms and test if a drug candidate could inhibit the virus-caused CVD or facilitate the disease recovery(23). However, heterozygous *Ifnar1^+/−^*and *Ifnag^+/−^* mice survive CHIKV infection and exhibit the highest viral presence in the heart on D1, followed by a rapid decline from D2 to D4, and then became very low, even undetectable, by D5 p.i. Immunohistochemistry and flow cytometry revealed that more leukocytes, particularly neutrophils, infiltrated the hearts of *Ifnag^+/−^* and *Ifnar1^+/−^* mice than in WT mice(23). In addition, the H&E staining analysis showed that CHIKV infection caused vasculitis in the left ventricles on D5 p.i. in both heterozygous groups. These results suggest that the heterozygous *Ifnar1^+/−^* and *Ifnag^+/−^*mice are invaluable for studying pathogenesis and testing therapeutic interventions for CHIKV-caused cardiac diseases. Importantly, these mice permit viral replication in the cardiac tissues while preserving sufficient immune competence to interrogate immunoregulatory mechanisms, including IL-17A signaling. A previous study reported that innate immune signaling via the mitochondrial antiviral signaling protein (MAVS) is required for CHIKV clearance from the heart(72). Mice deficient in MAVS signaling show persistent infection, leading to focal myocarditis and vasculitis of the large vessels attached to the base of the heart(72). In contrast, *Ifnag^+/−^* mice survive CHIKV infection while developing significant cardiac inflammation, including vasculitis, leukocyte and neutrophil infiltration, and early profibrotic responses. Therefore, we employed the *Ifnag^+/−^* mouse model to examine the contribution of IL-17A signaling to CHIKV-induced cardiac injury.

We observed that CHIKV infection upregulates *Il-17a* expression in the heart; thus, we assessed the role of IL-17A signaling in the pathogenesis of CHIKV-induced CVD by neutralizing IL-17A with α-IL-17A mAb. In the pretreatment experiments, blocking IL-17A signaling not only reduced viral burden in the heart and circulation but also significantly decreased the expression of proinflammatory cytokines and chemokines, including *Il-1β*, *Tnf-α*, and *Cxcl2.* Although tissue inflammation is one of the consequences of viral infection, excessive and prolonged inflammation is often more damaging than viral replication itself, leading to cardiomyocyte hypertrophy and reduced contractility (34, 35). The histopathological analysis confirmed that IL-17A neutralization preserved cardiac architecture in the left atrium and left ventricle, the most susceptible regions of the heart to CHIKV infection, by preventing dense immune cell infiltration. Flow cytometric analysis revealed that IL-17A is a primary driver of neutrophil and leukocyte recruitment to the heart. While neutrophils are essential for early defense, their excessive recruitment in isotype Ab-treated controls, compared with α-IL17A-treated mice, contributes to tissue remodeling and damage through the release of reactive oxygen species and proteases(36, 37). By limiting immune cell infiltration, α-IL17A treatment effectively prevents pathological inflammation and cardiac injury. Reduced cardiac injury following IL-17A signaling blockage was also confirmed by decreased levels of cardiac injury biomarkers, including troponin I, endocan, and connective tissue growth factor (CTGF). The elevation of Troponin I and Endocan in the isotype control groups mirrors clinical observations in fatal human CHIKV cases(40), where endothelial dysfunction and myocardial injury are prominent features. Furthermore, the reduction in *Ctgf* expression suggests that targeting IL-17A may prevent the early transitions toward cardiac fibrosis, a common long-term complication of viral myocarditis. Our post-infection treatment study further demonstrates that therapeutic neutralization of IL-17A effectively limits viral dissemination in the circulation and heart and reduces CHIKV-associated cardiac tissue damage, indicating the therapeutic potential of blocking IL-17A signaling to treat CHIKV-induced heart diseases.

Finally, our experiments in primary human cardiac fibroblasts and HEK 293 cells confirm the findings in the mouse model that IL-17A signaling enhances CHIKV infection in human cardiac and non-cardiac cells by inhibiting type I interferon responses. These results indicate great therapeutic potential for CHIKV-induced CVDs by pharmaceutically blocking IL-17A signaling. Consistent with our findings, monoclonal antibody blockade of the IL-17A pathway has successfully mitigated viral myocarditis in Coxsackievirus B3 models (73, 74). Therapeutic strategies targeting IL-17A signaling primarily use monoclonal antibodies, classified as biological disease-modifying antirheumatic drugs (bDMARDs), to treat autoimmune and inflammatory conditions driven by IL-17A (42). Current clinical approaches include direct inhibition of IL-17A using FDA-approved agents such as Secukinumab and Ixekizumab, or blocking the IL-17RA receptor with Brodalumab(42). For instance, Brodalumab (AMG 827), a human monoclonal antibody with high affinity for IL-17RA, is used to treat plaque psoriasis(75). In this study, we revealed that Brodalumab effectively blocks IL-17A-induced CHIKV replication in human cardiac fibroblasts and HEK 293 cells, suggesting the feasibility of repurposing IL-17A signaling inhibitors, including Brodalumab, for CHIKV diseases, including cardiac complications.

## Conclusion

Our study identifies the IL-17A/IL-17RA signaling axis as a key driver of CHIKV-induced cardiac pathology. Inhibition of IL-17A enhanced type I interferon responses, reduced viral load, and limited immune cell infiltration, resulting in significant attenuation of cardiac injury in both murine models and human cells. In addition, our results lay the foundation for several important directions in future research on viral myocarditis. Although the present study establishes the IL-17A/IL-17RA axis as a central contributor to cardiac inflammation, the downstream molecular pathways through which IL-17A promotes tissue injury remain to be fully elucidated. Further investigation of these mechanisms may reveal additional therapeutic targets and provide greater insight into the regulation of antiviral and inflammatory responses during infection. Moreover, extending these studies to other alphaviruses will help determine whether IL-17A-driven inflammation represents a conserved pathogenic mechanism across related viral infections. Such work could support the development of broad-spectrum immunomodulatory strategies for the treatment and prevention of viral-induced cardiac diseases.

## Acknowledgements

The authors thank Dr. Robert B. Tesh (University of Texas Medical Branch) for providing CHIKV strain LR OPY1 2006, Dr. Richard A. Flavell (Yale University School of Medicine) and Dr. Sarah Gaffen (University of Pittsburgh) for providing the breeding pairs of *Il17a^−/−^* and *Il17ra^−/−^*mice, respectively. The authors also thank the Mississippi INBRE (funded by the National Institute of General Medical Sciences GM103476) for use of the research facility. This work was supported by the American Heart Association grants (26BAIREA1560965 and 23AIREA1051360) and the National Institutes of Health grant (R15AI178654).

## Author Contributions

F.B. conceived the experiments; S.U.K. conducted the experiments and analyzed the data; P.M.D. D., S.S., A.O., and N.S.B assisted in the experiments; F.B. and S.U.K. wrote the manuscript. All authors read and approved the manuscript.

## Data Availability

All the data will be available upon request.

## MATERIALS AND METHODS

### Ethics Statement and Biosafety

All experiments involving the live chikungunya virus (CHIKV) were conducted by certified personnel within the USDA-certified Biosafety Level 3 (BSL3) facility at the University of Southern Mississippi (USM), in accordance with the USM Institutional Biosafety Committee approval (protocol #20250207). Animal procedures were approved by the USM Institutional Animal Care and Use Committee (IACUC; protocol #15101601). Mice were anesthetized using 25% v/v isoflurane and humanely euthanized via carbon dioxide inhalation, adhering to the American Veterinary Medical Association (AVMA) Guidelines for the Euthanasia of Animals.

### Viruses and Cells

The CHIKV LR OPY1 2006 strain was provided by the World Reference Center for Emerging Viruses and Arboviruses at the University of Texas Medical Branch. A viral stock was generated via a single passage of the parental virus in Vero cells (ATCC CCL-81). Viral titers were determined by plaque-forming assay on Vero cells as previously described(20, 76). Vero and Human Embryonic Kidney (HEK 293) cells were maintained in Dulbecco’s Modified Eagle Medium (Gibco™ DMEM, Life Technologies) supplemented with 10% fetal bovine serum (FBS), 1% L-glutamine, and 1% penicillin/ streptomycin. Human cardiac fibroblasts (HCFs; Innoprot) were cultured in Fibroblast Medium-2 (Innoprot) containing 5% FBS, 1% fibroblast growth supplement 2, and 1% penicillin/streptomycin.

### Mice and Animal Study

Breeding pairs of wild-type (WT) C57BL/6J mice (Strain #: 000664), and *Ifnag ^−/−^* mice on a C57BL/6J background (Strain #:029098) were obtained from The Jackson Laboratory (Bar Harbor, ME). Heterozygous *ifnag^+/−^* mice were generated by mating the homozygous mice with WT mice. *Il17a^−/−^* and *Il17ra^−/−^* mice breeding pairs on a C57BL/6J background were generous gifts from Dr. Richard A. Flavell (Yale University School of Medicine) and Dr. Sarah Gaffen (University of Pittsburgh), respectively. Animals were housed in a clean facility, while all live CHIKV infection experiments were performed in an ABSL-3 laboratory. Mice received a subcutaneous injection of 1×10^5^ PFU of CHIKV in phosphate-buffered saline (PBS) into the ventral side of the left hind footpad. Blood was drawn via the retro-orbital sinus. Following anesthesia and subsequent systemic perfusion with cold PBS, hearts were collected for RT-qPCR, plaque-forming assays, flow cytometry, and immunohistochemistry. Primary cardiac fibroblasts were isolated from uninfected mice, and hearts were collected, minced, and enzymatically digested using a collagenase-DNase cocktail following a previous article(77).

Fibroblasts were isolated by selective adhesion and cultured in 6-well plates for CHIKV infection. For IL-17A-neutralization experiments, mice received intraperitoneal injections of either an anti-IL-17A antibody (100 μg/mouse, BioXCell; clone 17F3) or an isotype control (IgG, clone MOPC-21, 100 μg/mouse, BioXCell) on D-1, D0, or 6 hours post-CHIKV infection.

### Reverse Transcription-Quantitative PCR (RT-qPCR)

Total RNA was extracted from blood and heart tissues using TRIzol-Reagent (Invitrogen). First-strand complementary DNA (cDNA) was subsequently synthesized using the iScript cDNA Synthesis Kit (Bio-Rad). RT-qPCR assays were executed on a CFX Connect Real-Time System (Bio-Rad) using iTaq Universal Probes Supermix (Bio-Rad) to detect *CHIKV-E1* and mouse *β-actin*. Viral RNA copy numbers were calculated or expressed as the ratio of *CHIKV-E1* to *β-actin*. Relative fold changes (RFC) compared to the control group were determined using the comparative threshold cycle ^ΔΔ^CT after normalization to cellular *β-actin*. A comprehensive list of all primers utilized in this study is provided in Table 1.

**Table 1:**
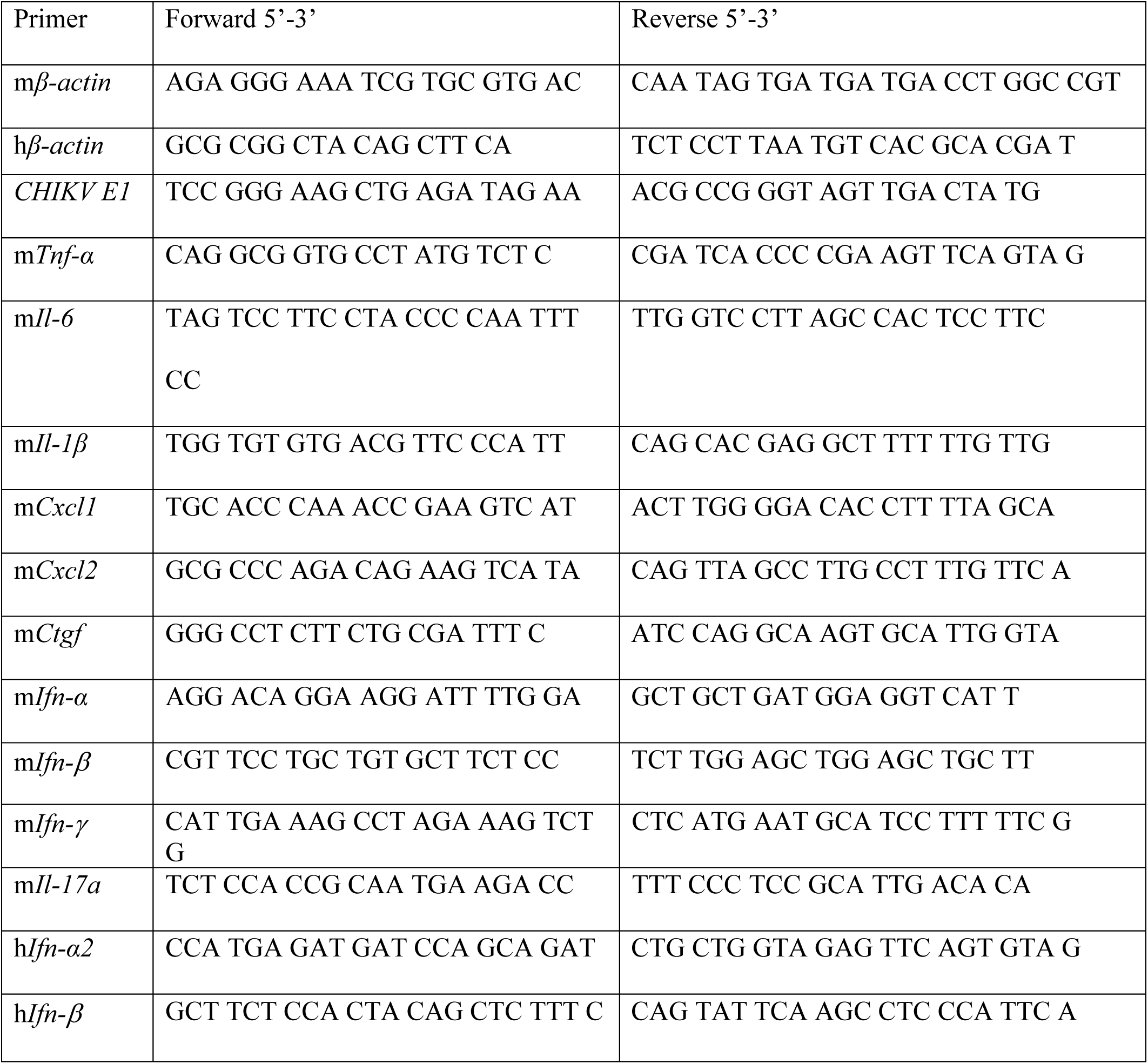
Primer sequences used for RT-qPCR.

### Plaque-forming Assay

Vero cells were seeded in 6-well plates at a density of 6 × 10^5^ cells per well and incubated overnight. Following centrifugation of homogenized heart tissue samples, the collected supernatants were serially diluted and inoculated onto confluent Vero cell monolayers. After incubation of 1 hour at 37°C with 5% CO_2_, the virus inoculum was removed, and the monolayer was overlaid with 1% SeaPlaque agarose (Lonza). The plates were incubated at 37°C with 5% CO_2_ for 24-48 hours to allow plaque formation. Plaques were visualized by staining with Neutral Red for 3 hours and subsequently counted. The viral titers were calculated and expressed as plaque-forming units per milliliter (PFU/ml) using the formula described in previous studies (78–80).

### Flow Cytometry

To analyze cardiac-infiltrating leukocytes, 3-week-old *Ifnag^+/−^*mice were inoculated via footpad injection with 1 × 10^5^ PFU of CHIKV or PBS (mock control) via footpad. On D1 p.i., mice were euthanized and perfused, and hearts were harvested. The cardiac tissue was minced and digested for 45 minutes at 37°C in a collagenase cocktail containing DNase I (Thermo Scientific), HEPES (Gibco™), and collagenase (Sigma) on a rotating shaker. Post-digestion, the cellular suspension was centrifuged at 393 g for 5 minutes at 4°C. The supernatant was removed, and the remaining cell pellet was resuspended in 3 mL of Hank’s Balanced Salt Solution (HBSS, Gibco™) before being passed through a 40 µm cell strainer. For surface marker staining, isolated leukocytes were labeled with fluorochrome-conjugated antibodies against CD45 (PE-Cy7, eBioscience), CD11b (APC, BD Biosciences), and Ly6G (APC-Cy7, Invitrogen). Samples were acquired on a BD LSRFortessa™ Cell Analyzer (BD Biosciences) and analyzed using FlowJo software (v10.9.0).

### Immunofluorescence assays

*Mouse Heart:* Three-week-old *Ifnag^+/−^* mice were inoculated with 1 × 10^5^ PFU of CHIKV via footpad injection. On D1 p.i., mice were perfused, and hearts were harvested and fixed in 4% paraformaldehyde (PFA). Tissues were sequentially immersed in 20% and 30% sucrose solutions for 24 hours at 4°C to provide cryoprotection. The cryoprotected hearts were then oriented on Tissue-Tek trays (Sakura Finetek USA, Inc.), embedded in optimal cutting temperature (OCT) compound (Fisher HealthCare), and snap-frozen in liquid nitrogen prior to overnight storage at −80 °C. Longitudinal cardiac sections were cut at a thickness of 10 µm using a cryostat (Tissue-Tek Cryo3, Sakura), mounted onto charged glass slides (Fisher Scientific), and air-dried overnight. Sections were subsequently incubated with 200 µl of antigen retrieval solution and blocked for 1 hour at room temperature using 2% goat serum supplemented with 0.3% Triton X-100 and 0.05% Tween-20 in PBS. The sections were probed overnight at 4^°^C with the mouse monoclonal anti-Chikungunya E2 antibody (1:100, Leinco Technologies, Inc.). After washing 2 times with 0.1% PBST (PBS with 0.1% Tween 20) on a shaker, cells were stained with Alexa Fluor 488-conjugated anti-mouse IgG (1 µg/mL, Invitrogen) for 2 hours at 4^°^C. After three subsequent PBST washes, cells were stained with DAPI (300 nM, Invitrogen).

*Human cardiac fibroblasts (HCFs)*: HCFs were pre-treated with Brodalumab (Selleckchem), a human monoclonal antibody targeting the IL-17 receptor A (IL-17RA), at concentrations of 50 ng/mL and 100 ng/mL for 3 hours. Following pre-treatment, cells were organized into three experimental groups: (1) Brodalumab pre-treatment followed by CHIKV infection (MOI 0.1 or 0.5) and supplementation with 50 ng/mL human recombinant IL-17A (rIL-17A, R&D Systems); (2) CHIKV infection with rIL-17A supplementation in the absence of Brodalumab; and (3) CHIKV infection alone at the specified MOIs. On D1 p.i., cells were fixed with 4% PFA for 20 minutes, then permeabilized with 0.1% Triton X for 30 minutes, and blocked with 2% Bovine Serum Albumin (BSA) containing 0.5% Triton X-100 and 0.05% Tween-20 in PBS for 1 hour at room temperature. Then, cells were probed overnight at 4^°^C with the mouse monoclonal anti-Chikungunya E2 antibody (1:100, Leinco Technologies, Inc.). After washing 2 times with 0.1% PBST (PBS with 0.1% Tween 20) on a shaker, cells were stained with Alexa Fluor 488-conjugated anti-mouse IgG (1 µg/mL, Invitrogen) for 2 hours at 4^°^C. After three subsequent PBST washes, cells were stained with DAPI (300nM, Invitrogen). The images were captured using a Stellaris STED confocal microscope (Leica).

### Hematoxylin and Eosin Staining

Pathological examination of cardiac tissue was conducted using a Hematoxylin and Eosin (H&E) Staining Kit (Abcam). Heart samples embedded in OCT compound were sectioned longitudinally at a thickness of 10 µm. Sections were first hydrated in distilled water and then incubated in Hematoxylin (Modified Mayer’s Solution) for 5 minutes. Following two rinses in distilled water, slides were immersed in a bluing reagent for 15 seconds, washed in two changes of distilled water, and dipped briefly in 95% ethanol. Counterstaining was performed using Eosin Y Solution for 2 minutes. The tissue sections were then sequentially dehydrated through one change of 95% ethanol and three changes of 100% ethanol. Finally, the slides were cover-slipped using a mounting medium (Surgipath Micromount, Leica). This protocol was adapted from previously established methodologies(20, 81). Histological images were captured using a Nikon Eclipse 80i microscope (Nikon, Japan).

### Isolation of mouse cardiac fibroblasts

After euthanasia and cervical dislocation, mouse bodies were sprayed with 70% ethanol and placed ventral side up. The abdominal skin and muscle were cut open, and a vertical incision was made toward the sternum to expose the thorax. The ribs were carefully cut to reveal the heart using autoclaved forceps and scissors and rinsed in cold Kreb’s-Henseleit buffer (KHB) buffer to remove excess blood. Hearts were transferred to a 10 cm² plate with 1 mL of digestion cocktail and minced with a single-edge blade. An additional 1 mL digestion cocktail was added, and mincing continued until fragments were pipettable. Fragments were transferred to a 50 mL conical tube, and the plate was washed twice with 2 mL digestion cocktail. The tube was incubated at 37°C with agitation, and the tissue was resuspended 10 times with a 5 mL pipette after 15 minutes. After a second 15-minute incubation and resuspension, the digested material was diluted in 25 mL KHB, filtered through a 40 µm cell strainer, and centrifuged at 400 g, 4^°^C for 10 minutes. The pellet was treated with RBC lysis buffer (5 mL/heart) for 2 minutes at RT, centrifuged (400 g, RT, 10 minutes), and resuspended in 1 mL KHB. After adding 9 mL KHB and filtering again, the sample was centrifuged, and the final pellet was resuspended in 1 mL DMEM-F12. Cells were counted, and 1 mL of suspension was plated per well of a 6-well plate pre-filled with 2 mL fibroblast media. After 4 hours of incubation at 37 °C, unattached cells were removed, wells were washed with PBS, and replaced with 2∼4 mL fresh medium. Medium was refreshed every 2 days until confluence. 2 × 10⁵ cells/well were seeded in 12-well plates and infected with CHIKV at an MOI of 1.0 for 1 hour. After infection, the inoculum was removed, the wells were washed with PBS, and fresh medium was added. Cell culture medium was collected daily from D1 to D4 p.i. to determine viral titers by plaque-forming assay.

### Statistical analysis

Data were analyzed using the Mann-Whitney U test or two-tailed Student’s *t*-tests with GraphPad Prism (version 10.2.3), as appropriate.

